# Hamster (*Cricetus cricetus*) and ground squirrel (*Spermophilus citellus*) in the metropolitan area of Vienna, Austria: Urban development vs. species protection

**DOI:** 10.1101/2024.02.08.579520

**Authors:** Franz Rubel

**Affiliations:** Climate Change & Infectious Diseases Group, University of Veterinary Medicine Vienna, Austria

**Keywords:** Endangered species, Habitat loss, Eco-offsetting, Eco-compensation, Urban development

## Abstract

Although European hamsters and European ground squirrels are listed as critically endangered on the *IUCN Red List of Threatened Species*, their habitats are becoming increasingly smaller due to the rapid expansion of the city of Vienna. The distribution of both species over the period 2000–2023 was documented by compiling a map based on georeferenced data from scientific monitoring and citizen science collections. Based on this, the population sizes in the metropolitan area of Vienna were estimated at 4000 hamsters and 16000 ground squirrels. An overview map was also created for each of the two species, which contains the names of the most important local occurrences. Satellite images were used to document populations that became extinct during the study period or in which animals lost large parts of their traditional habitat due to new urban development. This is considered particularly important in terms of conservation, as future generations would otherwise see the absence of wildlife in cities as normal, an effect known as shifting baseline syndrome. The Court of Justice of the European Union clarified key concepts for strict species protection. These also include eco-offsetting, i.e. creating habitats to replace ones lost to development, which was already applied before a recent construction project was implemented. On the occasion of this project, the application of eco-offsetting to protect the local ground squirrel population was documented and critically discussed.

## 1. Introduction

The Common hamsters (*Cricetus cricetus* Linnaeus, 1758), also known as European hamster, and the European ground squirrel (*Spermophilus citellus* Linnaeus, 1766) are rodents that were considered agricultural pests for centuries. Today they are considered endangered species and are strictly protected. Within the subfamily Cricetinae, *C. cricetus* (Fig. 1, left) is the largest specimen with a body length of 200–300 mm and a body mass of 200–650 g, although animals with a body mass of 1000 g have also been described (Weinhold, 2008). Specimens of *S. citellus* (Fig. 1, right) are on average slightly smaller and lighter with a length of 18–23 cm and a body mass of 130–380 g. *Cricetus cricetus* is a nocturnal or crepuscular rodent that lives singly, though in a complex burrow system. The fertile steppe, forest steppe and grassland are its original habitats, where in some regions they occur sympatric with *S. citellus*. This is particularly true for the metropolitan area of Vienna considered here. *Spermophilus citellus* is also a rodent that lives in burrow systems, but is diurnal and organized in colonies. Both species have successfully spread into a variety of anthropogenic habitats, including cities (C^̌^ańady, 2013; Surov et al., 2019). For example, *C. cricetus* also occurs on pastures, arable land, field edges, fallow bushes, but also on roadsides and railway embankments (Weinhold, 2008). Human-made habitats include gardens, orchards, and habitats in urban environments such as cemeteries (Feoktistova et al., 2013) or green spaces between multi-storey residential buildings (Franceschini and Millesi, 2001; Flamand et al., 2019). In these habitats, weeds are the predominant food source (Tissier et al., 2019). *Spermophilus citellus* also occurs in maintained orchards, vineyards, and on wasteland between industrial plants (Hoffmann et al., 2008; Poledńıková et al., 2022).

**Figure 1:**
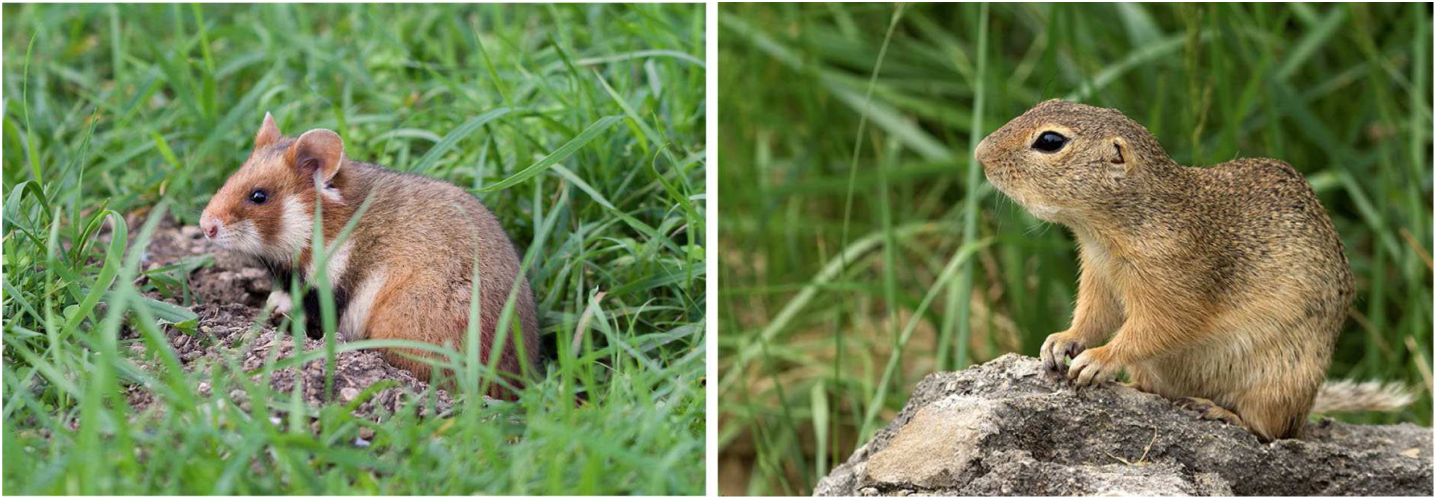
Common hamster (*Cricetus cricetus*) with a length of around 20–30 cm and a body mass of around 200–650 g (left) and European ground squirrel (*Spermophilus citellus*) with a length of 18–23 cm and a body mass of 130–380 g (right).

The global distribution of *C. cricetus* originally extended from Belgium in the west via the steppes of Kazakhstan to the Russian Altai Mountains in the east, within the range 43–58^◦^ N and 6–93^◦^ E (Surov et al., 2016; Feoktistova et al., 2017). The occurrence of *C. cricetus* has been documented in Austria, Belarus, Belgium; Bulgaria, Czechia, France, Germany, Hungary, Kazakhstan, Moldova, Netherlands, Poland, Romania, Russian Federation, Serbia, Slovakia, Slovenia, Switzerland, and Ukraine. Over the last few decades, a sharp decline in the *C. cricetus* population of approximately 75% has been observed (Surov et al., 2016), as a comparison of the distribution maps from 1970 to 2010 shows (Tissier et al., 2019). Simulation studies indicate that intensified farming practices in Europe are responsible for this (La Haye et al., 2014). However, in large parts of the Russian distribution area, the Caucasus countries and Crimea, the hamster population is still classified as consistently high (Bogomolov et al., 2022). The global distribution of *S. citellus* can be seen, for example, from the map by Ramos-Lara et al. (2014). Accordingly, *S. citellus* is today endemic to Central and Southeastern Europe, within the range 40–51^◦^ N and 13–29^◦^ E. The western distribution limit is in the Czech Republic and the eastern limit is on the European Black Sea coast of Turkey, whereby *S. citellus* has also been documented in Austria, Bulgaria, Greece, Hungary, Moldova, North Macedonia, Poland, Romania, Serbia, Slovakia and Ukraine. The distribution area of *S. citellus* is therefore significantly smaller than that of *C. cricetus*, but overlaps with them in parts of Central Europe and the Balkans. Like the European population of *C. cricetus*, the population of *S. citellus* is also declining rapidly and is already extinct in Germany and Croatia. Only in Bulgaria there are still some abundant populations of *S. citellus* (Koshev et al., 2023).

Both species, *C. cricetus* (Banaszek et al., 2020) and *S. citellus* (Hegyeli, 2020), have most recently been assessed for *The IUCN Red List of Threatened Species* in 2019 where they were listed as *Critically Endangered* under criteria A3c. In Austria, the two species are only endemic in the northeast of the country, with the largest urban *C. cricetus* population in the capital Vienna (Feoktistova et al., 2013).

This paper provides the international scientific community with an overview map and several detailed maps of the distribution of *C. cricetus* and *S. citellus* in Vienna, based on several local project reports published in German language and citizen science data. The maps document the distribution of the two species from 2000 to the present, whereby extinct populations and populations that were relocated as a result of urban expansion were also mapped. The maps can therefore be used at a later date to assess the development of the two strictly protected species, particularly with regard to the conflict between species protection and urban expansion. Finally, the distribution maps allow indirect conclusions to be drawn about the occurrence of specific parasites for which there is no current data. An example of the latter is the tick *Ixodes laguri*, which is considered a vector of *Francisella tularensis*. Typically, it infests burrowing rodents, mainly Cricetidae, Sciuridae and Gliridae and is co-distributed with them (Mihalca and D’Amico, 2017).

## 2. Materials and methods

Vienna, the capital of Austria, is located at the geographical coordinates 48.21^◦^ N/16.37^◦^ E. As of the beginning of 2024, it has approximately two million inhabitants, and covers an area of 415 km^2^ of which approximately 50% are green spaces (forest, agricultural land, parks, gardens and nature reserves) and water areas (Danube with tributaries). The city of Vienna is surrounded by the Province of Lower Austria, whose adjacent regions are counted to the metropolitan area of Vienna. It is characterized by a warm temperate climate with precipitation in all seasons and hot summers, which is classified after Köppen-Geiger as a Cfa climate. As a result of global warming, the previously warm summers (Cfb) have been replaced by hot summers (Cfa) in the last decade (Rubel et al., 2017). Fig. 2 shows the seasonal cycles of the monthly temperature and precipitation observations in Vienna for the period 1996–2020 with an annual mean temperature of 11.4^◦^C and an accumulated precipitation of 664 mm/year.

**Figure 2:**
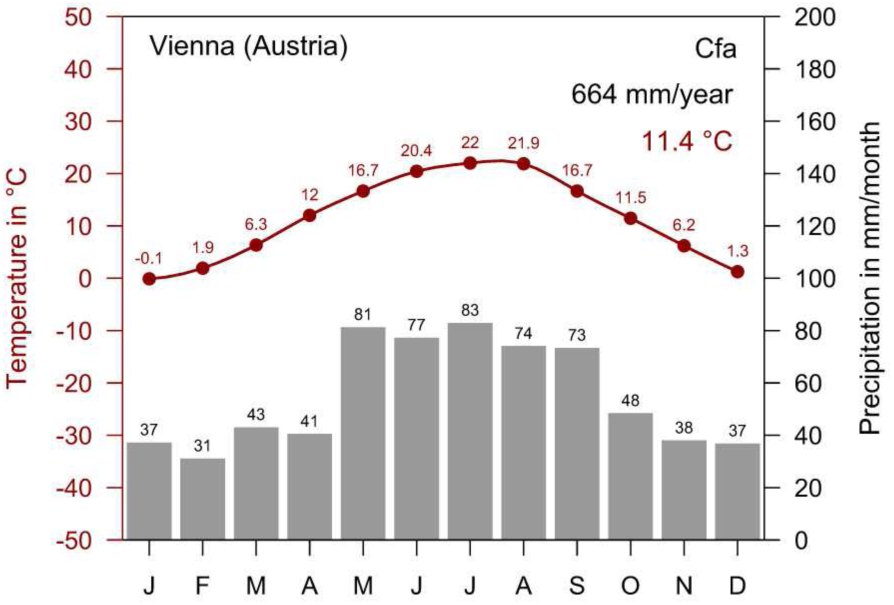
Climate diagram representative for the metropolitan area of Vienna, Austria. Climate classification after Köppen-Geiger Cfa, annual mean temperature 11.4^◦^C, accumulated precipitation 664 mm/year (period 1996–2020).

Mapped locations of *C. cricetus* and *S. citellus* from field studies mainly commissioned by the municipal government of the City of Vienna and the Province of Lower Austria were digitized and supplemented by locations provided by the Global Biodiversity Information Facility (GBIF). For this purpose, all locations of *C. cricetus* and *S. citellus* were extracted from the GBIF database. However, only data from the period 2000–2023, which is within the study region was used. This GBIF data also include locations from iNaturalist (iNaturalist Research-grade Observations), naturgucker (a German social network for nature watchers), HNS (Haus der Natur Salzburg, biodiversity database), and BOKU (data uploaded by the University of Natural Resources and Life Sciences, Vienna). All GBIF data are referred to below as citizen science data.

A total of 1252 georeferenced *C. cricetus* locations comes from six project reports of scientific monitoring programs (Table A.1). In this context, locations include sightings, active and inactive burrows and indirect evidence that indicate the occurrence of the species. Georeferenced locations are therefore not a direct measure of animal abundance, but are only intended to be used to visualize the distribution of the species. These data were supplemented by 208 citizen science locations (GBIF.org, 2023a) covering the period 2014–2023, which are exclusively sightings including 93 iNaturalist, 9 HNS, and 2 BOKU locations. The georeferenced *S. citellus* locations digitized for this study includes 1398 locations from nine project reports of scientific monitoring programs (Table A.2), which were supplemented by 90 citizen science locations (GBIF.org, 2023b). This citizen science dataset covers the period 2007–2023 and includes 43 iNaturalist, 3 naturgucker, 5 BOKU, and 2 HNS locations.

Maps were compiled with the R software (R Core Team, 2023) using the OpenStreetMap data from Stamen (https://stamen.com) and the virtual globe Google Earth (https://www.google.com/intl/en/earth). Overlapping data points have been filtered for clarity using a random selection and a thinning algorithm (Aiello-Lammens et al., 2015, 2019). For example, numerous sightings of *S. citellus* have been reported in the Perchtoldsdorfer Heide nature reserve in the southwest of the city of Vienna. However, only 3 out of 39 digitized locations were shown in the city map (Fig. 3) to avoid too much overlap of location points. The distance between the drawn points in the map of the metropolitan area of Vienna was set to 200 m. In contrast, all georeferenced locations are depicted on the high-resolution Google Earth maps.

**Figure 3:**
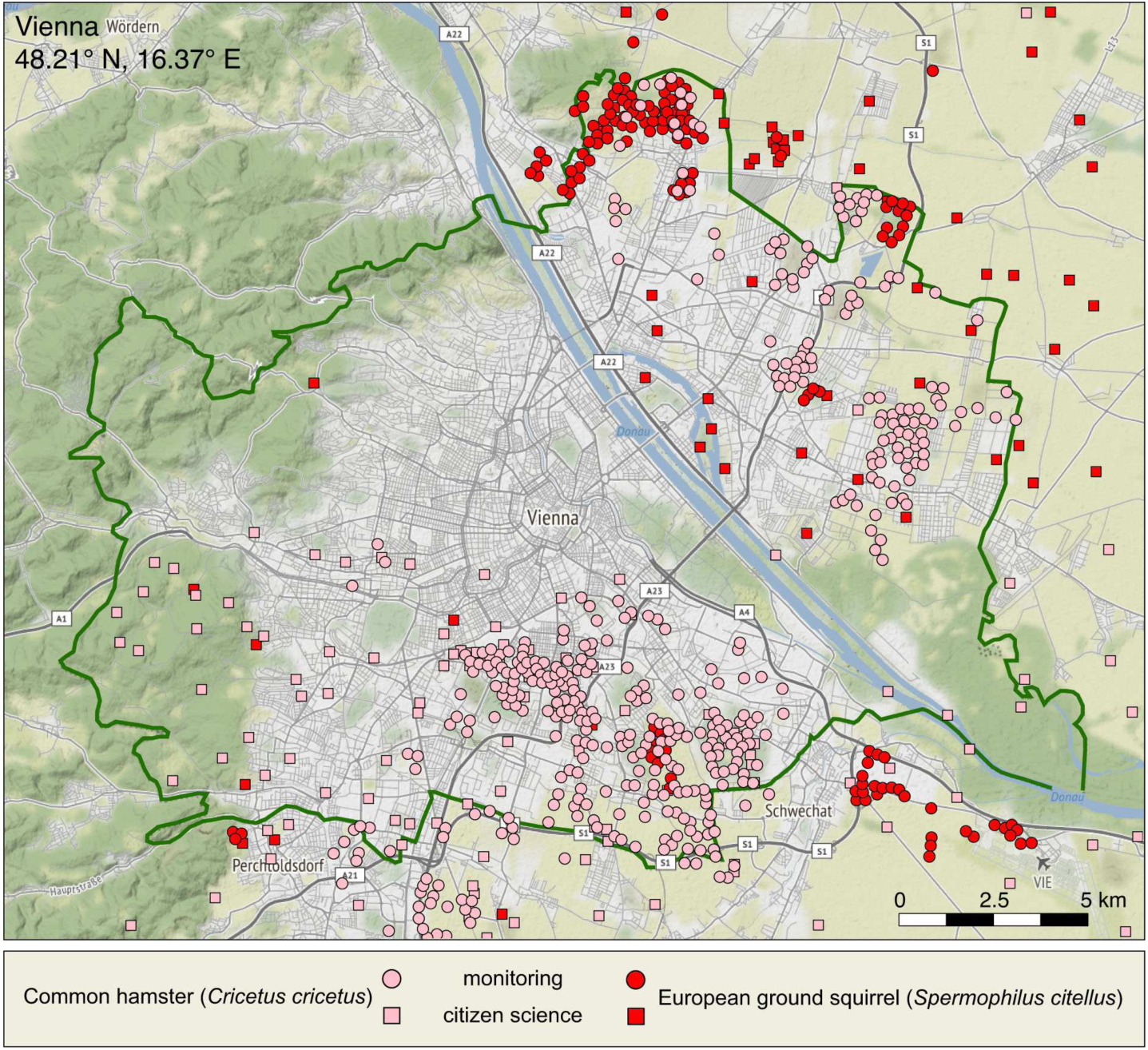
Recorded locations of *Cricetus cricetus* (pink) and *Spermophilus citellus* (red) in the metropolitan area of Vienna, Austria. Data from monitoring (circles) and citizen science collections (rectangles) are representative for the period 2000–2023.

## 3. Results and discussion

### 3.1. Mapped distribution and population sizes

An important result of this study is the distribution map of *C. cricetus* und *S. citellus* for the entire metropolitan area of Vienna, representative for the period 2000–2023 (Fig. 3). In general, it can be assumed that the georeferenced locations of these two species are more reliable if they come from scientific monitoring programs or from scientifically supported nature conservation projects. These locations were therefore represented on the map by circles, while mostly unconfirmed locations from citizen science data sets were represented by rectangles (Fig. 3). Citizen science data is mainly based on observations by laypeople and is considered less reliable. Nevertheless, these data are valuable because they cover otherwise unsampled regions, allowing the compilation of a more complete distribution map. In addition, citizen science data also contains observations from scientists such as those from the University of Natural Resources and Life Sciences mentioned above.

The mean population size of *C. cricetus* in the monitored regions of Vienna was estimated at 2400 individuals, with another 1000 individuals living in the central cemetery (Table A.3). If one takes into account the unsampled regions, then the hamster population in Vienna can be roughly estimated at 3600 individuals, which is the largest urban hamster population in Europe (Feoktistova et al., 2013). When one looks at the metropolitan area of Vienna, which also includes the outskirts of the city, the hamster population increases to an estimated 4000 individuals (Table A.3). The observations of *C. cricetus* mapped here (Fig. 3) show that with the exception of the inner city (districts 1, 2, 4–9) and the adjacent west of Vienna (districts 16–20), hamsters are documented throughout the entire city. Fig. 4(left) shows an overview map with the districts of Vienna and the largest occurrences of *C. cricetus*. The latter are north of the Danube in the 21st district (located at Stammersdorf and Marchfeld Canal, 900 animals) and in the 22nd district (Süßenbrunn, Kagran, Hirschstetten and Aspern, 400 animals). South of the Danube, the largest populations are documented in the 10th district (Favoriten with Laaerberg and Oberlaa, 750 animals), in the 11th district (Central Cemetery, 1000 animals), and in the 12th district (Meidling Cemetery, 150 animals). In Favoriten, the *C. cricetus* population on the grounds of the clinic Favoriten, the former Kaiser-Franz-Josef hospital, is particularly well documented (Flamand et al., 2019). The population sizes mentioned (Table A.3) are only rough estimates because the hamster populations are subject to annual fluctuations and developed differently from region to region during the period under consideration 2000-2023. Numerous citizen science observations in the southwest of Vienna indicate that there are *C. cricetus* populations in these districts that have not yet been recorded by monitoring programs. Outside the city limits, larger *C. cricetus* populations are mapped in the south of Vienna, in the so-called Thermal Region, which actually also includes Perchtoldsdorf.

**Figure 4:**
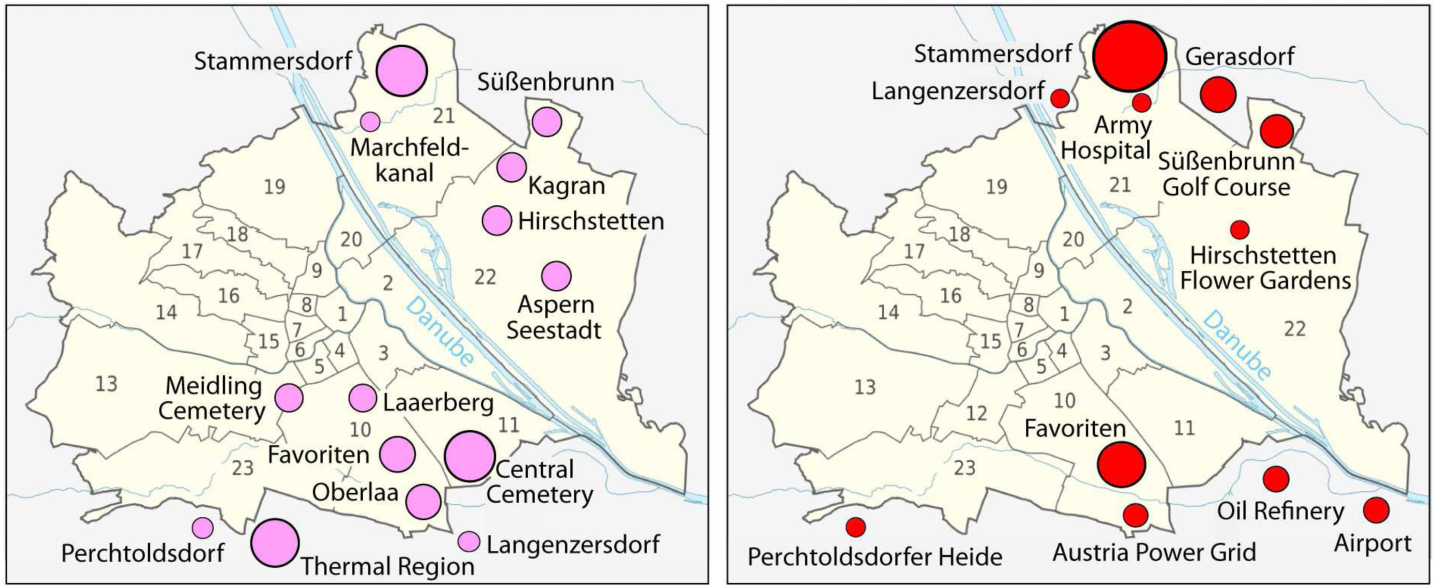
Overview of the main populations of *Cricetus cricetus* (left, pink) and *Spermophilus citellus* (right, red) in the metropolitan area of Vienna, Austria. Period 2000–2023.

The mean population size of *S. citellus* in the city of Vienna, based on the monitoring of 2020, was estimated at 14000 individuals. For the entire metropolitan area of Vienna including the outskirts, this estimate increases to 16000 individuals (Table A.4). The observations of *S. citellus* mapped here (Fig. 3) show several larger clusters summarized in Fig. 4(right). North of the Danube, these are the populations in Stammersdorf (9200 animals), on the grounds of the Stammersdorf Army Hospital (700 animals) and at the Süßenbrunn Golf Course (950 animals), as well as the population in Gerasdorf outside the city limits (900 animals). In Fig. 4(right) also smaller populations receiving a lot of public attention, are shown. These are the *S. citellus* populations in Langenzersdorf (Seeschlacht recreation area) with the neighbouring population at the Strebersdorf campus of the Religious Education Academy of the Archdiocese of Vienna (Fig. 5), and on the grounds of the Hirschstetten Flower Gardens (100 animals). South of the Danube, the *S. citellus* populations are, with the exception of a few, unverified citizen science observations, restricted to the centre of the 10th district of Favoriten and the occurrence at the Austria Power Grid site. These populations were estimated to be 1950 animals (Table A.4). There are also larger populations outside the city limits on the grounds and in the area surrounding the Schwechat Oil Refinery and the Vienna International Airport. The *S. citellus* populations at these locations were estimated to be 350 animals. The isolated occurrence of *S. citellus* at the Perchtoldsdorfer Heide (Fig. 5) is home to only some dozen animals, but is highly valued by the local human population and managed by the Lower Austria Nature Conservation Association. An average of 24 animals were counted in this protected area during the period 2015-2017. In the period 2019–2021, 95 additional adult animals were relocated from Wiener Neustadt, 35 km away, to the Perchtoldsdorfer Heide to strengthen the population. Soft release methods, a sufficiently large number of released animals, long-term management and regular monitoring of the newly established population according to the suggestions of Maťejů et al. (2012) have contributed to the successful relocation of these ground squirrels.

**Figure 5:**
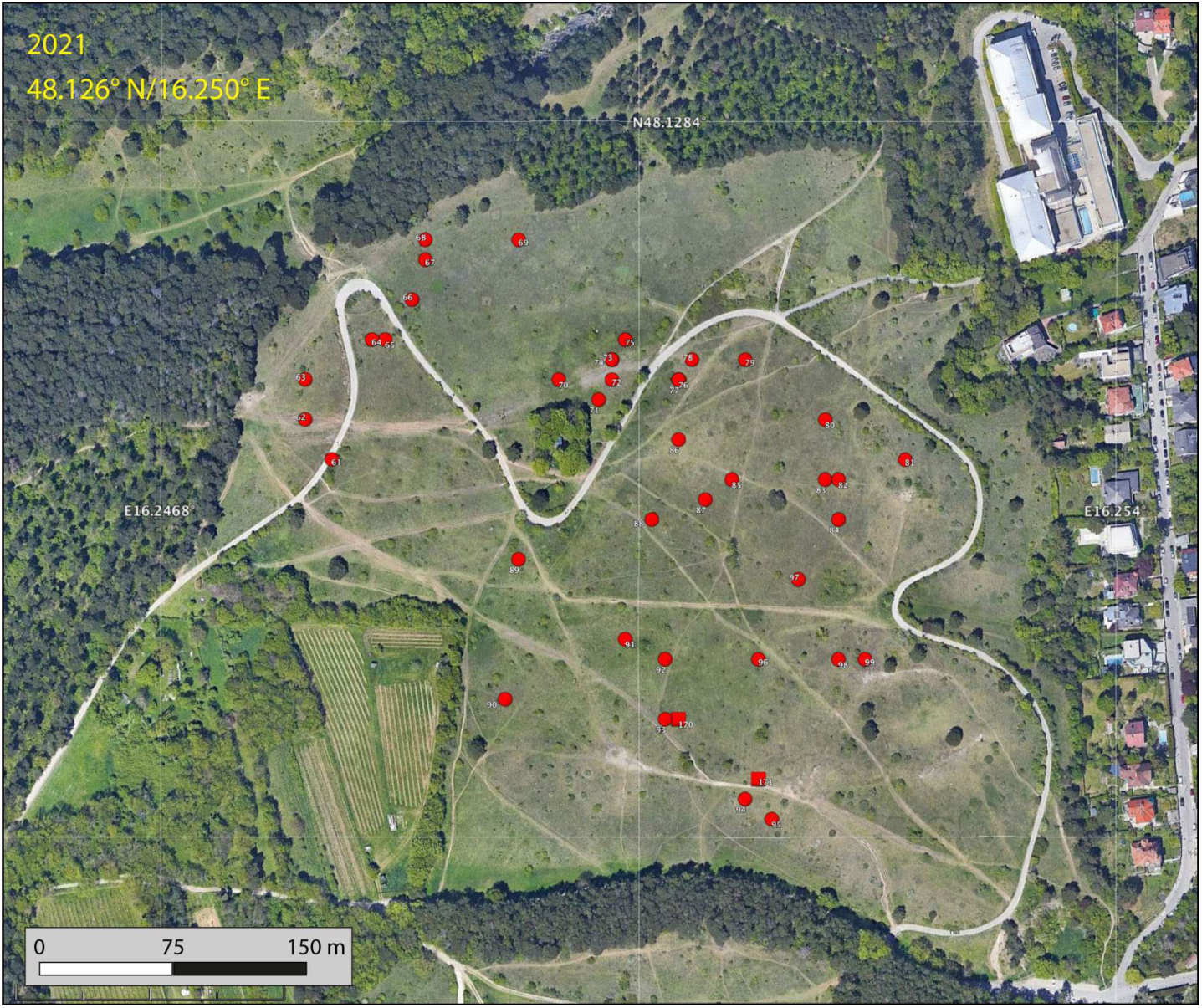
Isolated population of *Spermophilus citellus* at the Perchtoldsdorfer Heide, southwest of Vienna, depicted by Google Earth satellite images 5/2021. Data from monitoring 2017 (circles) and from citizen science collections (rectangles). In the period 2019-2021, animals from another population were released to strengthen the local population.

As mentioned above, the populations of both species fluctuate, sometimes considerably, during the period under consideration. This will be discussed below using selected examples.

### 3.2. Anthropogenically enforced population dynamics

Decreasing populations of *C. cricetus* and *S. citellus* are mainly due to the loss of suitable habitats. For example, the human population in Vienna increased from 1.55 to 1.98 million inhabitants during the period 2000–2023. This is associated with strong construction activity, particularly in the 21st and 22nd districts, which are considered urban expansion areas. Fig. 6 shows this development by comparing two Google Earth satellite images from the Aspern Seestadt. The satellite image from August 2014 (Fig. 6, above) shows the first construction phase on the former Aspern airfield surrounded by wasteland. The progress in the construction of residential buildings and the associated infrastructure can be clearly seen by comparing it with the satellite image from April 2021 (Fig. 6, below). *Cricetus cricetus* observations from the hamster monitoring 2015 (Table A.1) are superimposed and depict the loss of habitat, which can be expected to result in a significant decline in the *C. cricetus* population in this area once all work has been completed. The same applies to the extinction of the *S. citellus* population in Langenzersdorf, which lies in the commuter belt of the city of Vienna. The decline in *S. citellus* abundance in the so-called Seeschlacht recreation area (Fig 7, left) between 1992 and 1999 was analysed in detail by Hoffmann et al. (2003). This still intact but already significantly decimated *S. citellus* population, but also populations in and around the campus of the Religious Education Academy, was mapped in 2000. The last ground squirrel from the Seeschlacht recreational area was probably observed as part of the monitoring in 2006 (Table A.2). The construction of numerous terraced houses near the recreation area and the simultaneous conversion of adjacent fields into dense apple orchards were assumed as the main reasons for the extinction of the former large ground squirrel population. The *S. citellus* population is also threatened on the campus of the Religious Education Academy (Fig 7, right). Part of the area was already built in 2007 and a plan was discussed in 2021 that would allow additional 900–1000 apartments to be built on this site. However, the implementation of this construction project has been abandoned for the time being. Nevertheless, the ground squirrel and hamster populations north of the Danube will continue to be threatened in the future by new housing construction, which is rapidly reducing their potential available habitats.

**Figure 6:**
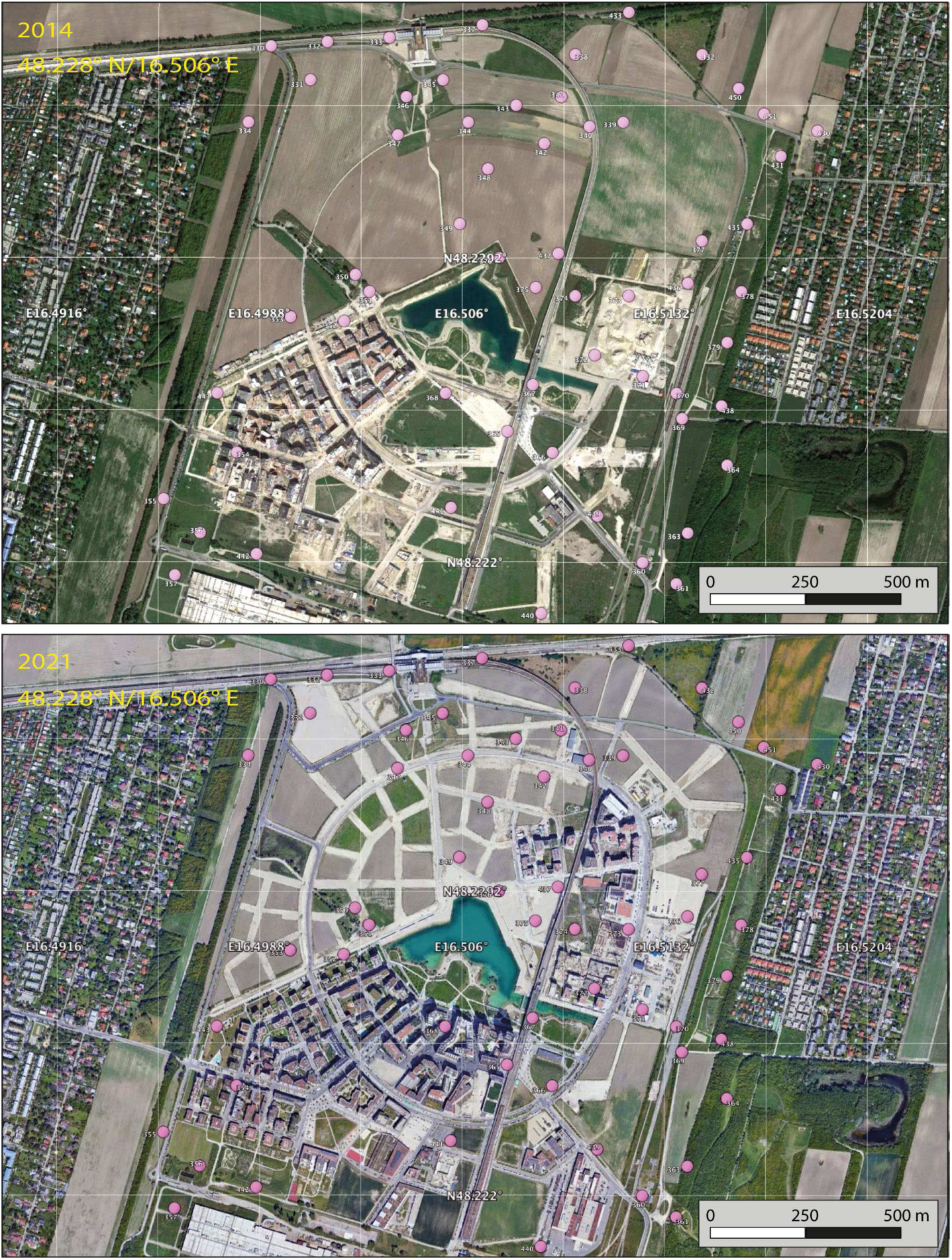
Habitat loss of *Cricetus cricetus* in Aspern Seestadt depicted by Google Earth satellite images 8/2014 (above) vs. 4/2021 (below). Data from monitoring 2015 (circles).

**Figure 7:**
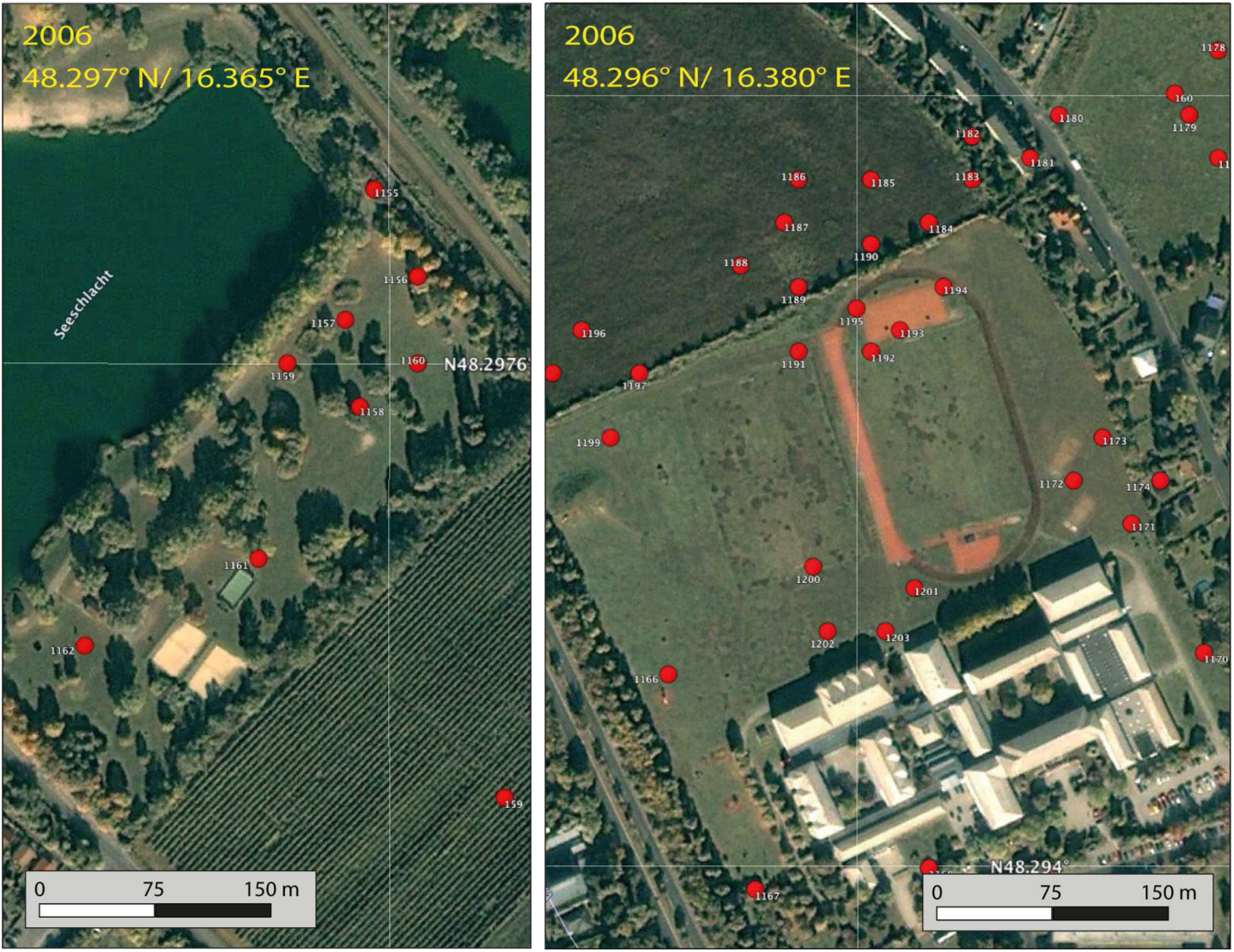
Populations of *Spermophilus citellus* in Langenzersdorf, Seeschlacht recreational area, extinct in 2007 (left) and on the neighboring campus of the Religious Education Academy, where a new building complex (not shown) was built north of the sports field in 2007. Current status of the population unknown (right). Google Earth satellite images from 10/2006 and data from monitoring 2000 (circles).

Increasing populations of *S. citellus* have also been reported. The estimated 14000 individuals of *S. citellus* in Vienna, mentioned above and based on the monitoring of 2020 (Table A.2), was reported to be higher than in previous years. An increase of *S. citellus* populations in the main occurrences of Stammersdorf, Süßenbrunn Golf Course, and Austria Power Grid is assumed to have been responsible for this. However, since new habitat areas of *S. citellus* have also been mapped, it has not been scientifically proven whether the propagated increase in the number of animals is real or can be attributed to different, temporally inhomogeneous monitoring strategies. Currently, the above-mentioned efforts to protect *S. citellus* in the Perchtoldsdorfer Heide also appear to be contributing to the stabilization of the ground squirrel population there.

### 3.3. Eco-offsetting – creating habitats to replace ones lost to development

Recently, the presence of the hamsters and ground squirrels north of the Stammersdorf Army Hospital (Fig. 4, right) interfered with future urban development plans, which presented the Court of Justice of the European Union (EU) with the opportunity to clarify certain key concepts that are entrenched in the system of strict species protection (Schoukens, 2022). The EU Habitats Directive for the strict protection of certain animal species does not differentiate between species found in rural and city habitats, or inside and outside protected areas. It also obliges Member States to put in place the necessary measures for the strict protection of these species, prohibiting all forms of deliberate capture or killing. Likewise, Member States must also prohibit all types of activities that lead to either the deterioration or destruction of the resting and breeding grounds of protected species. Furthermore, the analysis of López-Bao et al. (2018) shows that Articles 6 and 12 of the Habitats Directive (Directive 92/43/EEC) require Member States to restore populations that are quasi extinct. Therefore, legal construction measures that encroach on areas with protected species such as hamsters are generally prohibited. Only under restrictive conditions can a derogation be issued and the strict prohibitions be bypassed (Schoukens, 2022). In Vienna, interventions in the habitats of critically endangered species require special permits from the responsible authorities, which include remedial and compensatory measures. In practice, this means that construction projects may only begin after the affected animals have been relocated. To achieve this, suitable and protected replacement habitats must be made available to ensure the survival of the population. Such interference has to be accompanied by a person in charge of so-called ecological supervision (Hoffmann and Habert, 2023).

The change in the habitat of *C. cricetus* and *S. citellus* on the grounds north of the Stammersdorf Army Hospital demonstrates how this was implemented. The *S. citellus* sightings and burrows on and around the grounds of the Army Hospital before construction began in 2007 are shown in Fig. A.1. It can be seen that the area planned for development north of the Army Hospital (Fig. 8) is approximately a third of the distribution area of this population. The first satellite image from 8/2016 with overlaid monitoring data from 2015 (Fig. 8, above) shows the preparatory work of removing the upper 30 cm soil layer in order to get animals to leave their burrows before the construction begins. For this purpose, a so-called ground squirrel bridge was built in 2015, which is intended to enable the animals to colonize compensation areas east of the Marchfeld Canal. The second satellite image from 6/2023 with overlaid monitoring data from 2021 (Fig. 8, below) shows the completed construction projects. The monitoring commissioned by the City of Vienna (Magistrate Department 22 for Environmental Protection) in 2020/2021 demonstrates that the now smaller undeveloped area is still inhabited by both species, *C. cricetus* and *S. citellus*, and that the compensation areas offered are partially populated. Formally, the survival of the local hamster and ground squirrel populations can therefore be viewed as assured. However, the compensation areas offered east of the Marchfeld Canal were only sparsely occupied by the animals. Of the eight designated areas, only half were accepted by the animals. The monitoring reports show that 27 animals were documented on the compensation areas in 2015 (after the bridge was constructed and before construction of the residential complex began) and that a similar number was still reported in 2022. In contrast, there were 270 animals counted on the grounds north of the Stammersdorf Army Hospital in 2015, 113 in 2021, and 61 in 2022. This means that the hamster and ground squirrel population has decreased to less than a quarter. It can therefore be summarized that despite the provision of compensation areas, the animal population declined sharply after construction began.

**Figure 8:**
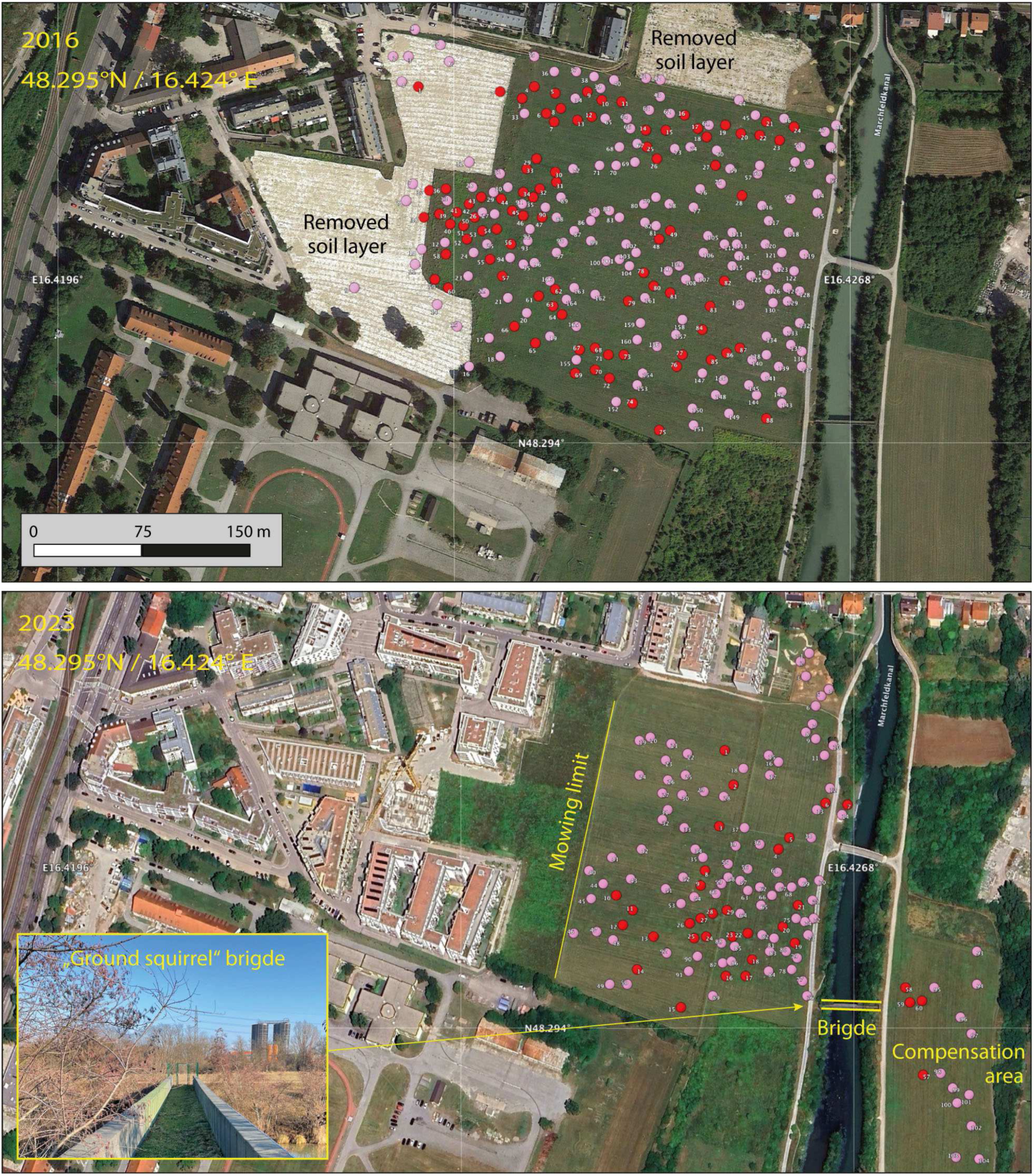
Locations of sightings as well as active and inactive burrows of *Cricetus cricetus* (pink, including unspecified burrows) and *Spermophilus citellus* (red) on the grounds north of the Stammersdorf Army Hospital. The Google Earth satellite image 8/2016 overlaid with monitoring data 2015 (above) shows the removal of the upper 30 cm soil layer to get animals to leave their burrows before construction begins. The recent satellite image 6/2023 overlaid with monitoring data 2021 (below) demonstrate the new housing construction and the so-called ground squirrel bridge to one of the compensation areas. Mowing limits the area reserved for hamsters and ground squirrels to the west (see also the entire hospital site in Fig. A.1).

The example shows that eco-offsetting is not an easy undertaking and also requires considerable effort. Ground squirrels in particular have special requirements concerning the height of the vegetation as well as the depth and proportion of sand and silt in the soil (Kenyeres et al., 2018), but also in terms of the dietary base for overwinter survival (Arok et al., 2021). The same applies to hamsters, which benefit from sown wildflower fields, for example (Fischer and Wagner, 2016). Obviously, the fragmented compensation areas are less suitable in this respect than the original areas. There is also a lot of use by dog owners who use the compensation areas to walk and exercise their dogs. This puts additional pressure on wildlife populations. In general, as urban development increases, the number of available compensation areas decreases and there is a risk that affected populations will become isolated. In addition, the pressure on wildlife is increasing, as the high number of new residents and dog owners lead to overuse of the remaining green spaces.

## 4. Conclusions

Although hamster and ground squirrel populations are declining almost throughout their entire range, they have adapted to semi-urban and urban habitats and reach a considerable number of estimated 4000 hamsters and 16000 ground squirrels in the metropolitan area of Vienna. In general, hamsters are found closer to the city centre than ground squirrels, which reach their largest population on the outskirts. South of the Danube, the largest hamster populations are in Favoriten, where the animals are found in parks and green spaces between residential buildings, as well as at the Central Cemetery and the Meidling Cemetery. There are also large populations of hamsters outside the city limits in the Thermal Region. North of the Danube, the Stammersdorf and Süßenbrunn populations are among the largest. Ground squirrels mainly inhabit semi-arid grassland, vineyards or alfalfa meadow on the northern outskirts in Stammersdorf, Gerasdorf and Süßenbrunn Golf Course and on the southern outskirts in Favoriten, on the grounds of the Austria Power Grid, of the Schwechat Oil Refinery and of the Vienna International Airport. In addition to a distribution map of the observations of both species (Fig. 3), an overview map was compiled from which the local occurrences mentioned can be seen (Fig. 4). The greatest danger to hamster and ground squirrel populations today comes from the rapid urban development north of the Danube. Several detailed maps document, among other things, the habitat loss of the *C. cricetus* population in Aspern Seestadt (Fig. 6) and the already extinct *S. citellus* population in Langenzersdorf Seeschlacht (Fig. 7) for new generations. By making information or experience with historical conditions available, the so-called shifting baseline syndrome is counteracted, according to which new generations accept the situation in which they were raised as normal. This contributes to the understanding that the presence of wild animals in urban environments is normal and therefore worth protecting.

Finally, eco-offsetting is currently the only compromise to reconcile urban development and species protection in Vienna. However, the example of the relocated animals on the grounds north of the Stammersdorf Army Hospital (Fig. 8) shows only limited success. In this urban development area, the hamster and ground squirrel population has decreased to less than a quarter. The present study demonstrates species conservation measures and may enable future assessments of whether hamster and ground squirrel populations have been successfully maintained in the long term.

## Acknowledgements

**Appendix A. Supplementary data**

