## Supplement for "Hamster (*Cricetus cricetus*) and ground squirrel (*Spermophilus citellus*) in the metropolitan area of Vienna, Austria: Urban development vs. species protection"

### Appendix A. Supplementary Material

The scientific monitoring data was only available in the form of maps that were published in German-language project reports. The field studies described in this grey literature were commissioned by authorities or required as part of building permits and carried out or at least accompanied by experts. Georeferenced locations include sightings as well as active and inactive burrows and are therefore not a direct measure of animal abundance. Unspecified hamster or ground squirrel burrows, e.g. by KNOLLCONSULT (2015, 2021, 2022), are listed in the Table A.1. Burrows clearly assigned to ground squirrels are listed in Table A.2.

Table A.1: Scientific monitoring reports of hamsters (*Cricetus cricetus*) in the metropolitan area of Vienna, provided by various clients and/or national authorities. The year of monitoring, the number of georeferenced locations, the figures in which they were used, and the references are given.

| Client/Authority | Year | No. | Fig. | Reference |
|---|---|---|---|---|
| City of Vienna, MA22 | 2010 | 50 | 3 | Hoffmann (2010) |
| LA Nat. Conserv. Assoc. | 2014 | 57 | 3 | Enzinger (2015) |
| Priv. property developers | 2015 | 166 | –, 8 | KNOLLCONSULT (2015) |
| City of Vienna, MA22 | 2015 | 451 | 3, 6 | Komposch et al. (2016) |
| City of Vienna, MA22 | 2020 | 411 | 3 | Hoffmann et al. (2020) |
| Priv. property developers | 2021 | 117 | –, 8 | KNOLLCONSULT (2021) |

Table A.2: Scientific monitoring reports of ground squirrels (*Spermophilus citellus*) in the metropolitan area of Vienna, provided by various clients and/or national authorities. The year of monitoring, the number of georeferenced locations, the figures in which they were used, and the references are given.

| Client/Authority | Year | No. | Fig. | Reference |
|---|---|---|---|---|
| City of Vienna, MA22 | 2000 | 71 | 3, 7 | Ulbel (2000) |
| City of Vienna, MA22 | 2011 | 353 | 3 | Hoffmann and Haberl (2011) |
| LA Nat. Conserv. Assoc. | 2013 | 58 | 3 | Enzinger (2014) |
| Priv. property developers | 2015 | 88 | –, 8 | KNOLLCONSULT (2015) |
| City of Vienna, MA22 | 2015 | 60 | 3 | Komposch et al. (2016) |
| Assoc. Friends PH | 2017 | 39 | 3, 5 | Drozdowski and Mrkvicka (2018) |
| LA Nat. Conserv. Assoc. | 2017 | 7 | 3 | Enzinger and Gross (2018) |
| City of Vienna, MA22 | 2020 | 662 | 3, 9 | Brunner et al. (2020) |
| Priv. property developers | 2021 | 60 | –, 8 | KNOLLCONSULT (2021) |

The following abbreviations were used: City of Vienna, MA22 (Magistrate Department 22 for Environmental Protection) = Stadt Wien, MA22 (Magistratsabteilung 22 für Umweltschutz); LA Nat. Conserv. Assoc. (Lower Austrian Nature Conservation Association) = NÖ Naturschutzbund; Assoc. Friends PH (Association of Friends of the Perchtoldsdorfer Heide) = Verein Freunde der Perchtoldsdorfer Heide, Priv. property developers = Private property developers or housing associations.

Table A.3: Estimated population size of hamster (*Cricetus cricetus*) in the metropolitan area of Vienna. Other regions refer to regions in which no scientific monitoring has been carried out, but where hamsters can be assumed to occur based on citizen science data (e.g. Western districts of Vienna, Danube-Auen National Park).

| Region | *C. cricetus* | Reference |
|---|---|---|
| City of Vienna | | |
| Stammersdorf & Marchfeld Canal | 900 | Komposch et al. (2016) |
| Süßenbr., Hirschst., Kagr., Aspern | 400 | Komposch et al. (2016) |
| Favoriten | 750 | Komposch et al. (2016) |
| Meidling, incl. Meidling Cemetery | 350 | Komposch et al. (2016) |
| Central Cemetery | 1000 | Hoffmann (2010) |
| Other regions | 200 | GBIF.org (2023a) |
| Provinz of Lower Austria | | |
| Thermal region south of Vienna | 300 | Enzinger (2015) |
| Other regions | 100 | GBIF.org (2023a) |
| **Total** | **4000** | |

Table A.4: Estimated population size of ground squirrels (*Spermophilus citellus*) in the metropolitan area of Vienna. Other regions refer to regions in which no scientific monitoring has been carried out, but where ground squirrels can be assumed to occur based on citizen science data.

| Region | *S. citellus* | Reference |
|---|---|---|
| City of Vienna | | |
| Stammersdorf | 9200 | Brunner et al. (2020) |
| Army Hospital | 700 | Hoffmann and Haberl (2011) |
| Süßenbrunn Golf Course | 950 | Brunner et al. (2020) |
| Gerasdorf | 900 | Brunner et al. (2020) |
| Favoriten & Austria Power Grid | 1950 | Brunner et al. (2020) |
| Hirschstetten Flower Gardens | 100 | Pledl (2010) |
| Other small occurrences | 200 | Brunner et al. (2020) |
| Provinz of Lower Austria | | |
| Oil Refinery & International Airport | 350 | Enzinger (2014) |
| Gerasdorf | 900 | Enzinger and Gross (2018) |
| Other regions | 750 | GBIF.org (2023b) |
| **Total** | **16000** | |

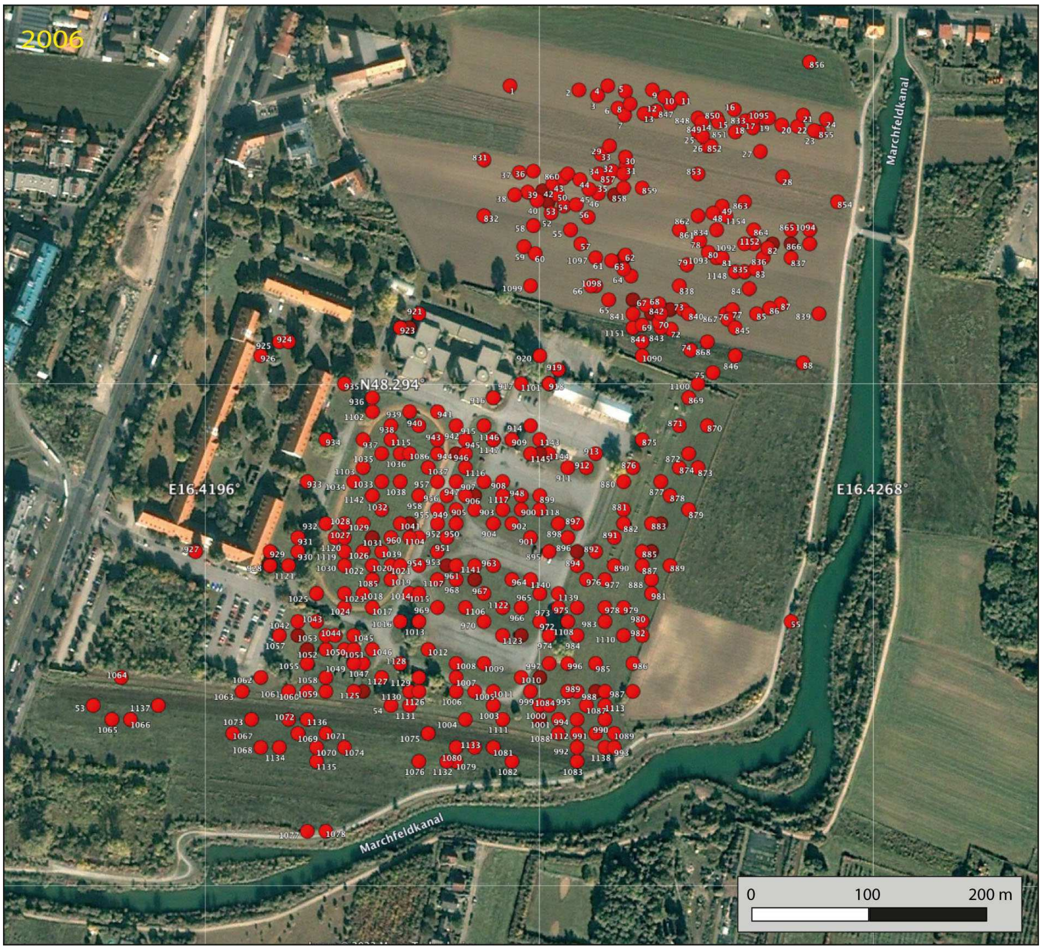


Figure A.1: Ground squirrel (*Spermophilus citellus*) sightings and burrows on and around the grounds of the Stammersdorf Army Hospital before construction began in 2007. Google Earth satellite images from 7/2006 and earliest monitoring data (circles) from Hoffmann and Haberl (2011) and KNOLLCONSULT (2015). The population was estimated at 600–850 ground squirrels and 55 hamsters (not shown) in 2010 (Hoffmann and Haberl, 2011).

KNOLLCONSULT, 2015. Bericht ökologische Aufsicht 14.07.2015. Projekt Knollconsult Umweltplanung ZT GmbH in Kooperation mit I. E. Hoffmann, Wohnbau nördlich Heeresspital – Zieselvorkommen, 8pp.

KNOLLCONSULT, 2021. Bericht ökologische Aufsicht 14.10.2021. Projekt Knollconsult Umweltplanung ZT GmbH in Kooperation mit I. E. Hoffmann, Wohnbau nördlich Heeresspital – Zieselvorkommen, 3pp.

KNOLLCONSULT, 2022. Bericht ökologische Aufsicht 12.10.2022. Projekt Knollconsult Umweltplanung ZT GmbH in Kooperation mit I. E. Hoffmann, Wohnbau nördlich Heeresspital – Zieselvorkommen, 4pp.

Komposch, B., Zimmermann, P., Holzinger, W., Hoffmann, I.E., 2016. Map of the co-occurrence of hamsters and ground squirrels in the city of Vienna. In: Bestandserfassung und -evaluierung des Feldhamsters (*Cricetus cricetus*) im Jahr 2015 im Stadtgebiet von Wien. Projektendbericht im Auftrag der Stadt Wien, MA22, 41pp. https://www.wien.gv.at/umweltschutz/naturschutz/biotop/feldhamster.html (accessed, 16 July 2023).

Pledl, G., 2010. Ziesel: Neue Großfamilie in den Blumengärten Hirschstetten zu bestaunen. Presseaussendung der Wiener Stadtgärten (MA 42), https://www.ots.at/presseaussendung/OTS_20100722_OTS0083 (accessed, 9 Jan 2024).

Ulbel, G., 2000. Gutachten zum Wiener Arten- und Lebensraumschutzprogramm Netzwerk Natur: Ziesel. Projektbericht im Auftrag der Stadt Wien, MA22, 6pp. https://www.zobodat.at (accessed, 16 July 2023).
